# The Arabidopsis receptor kinase STRUBBELIG regulates the response to cellulose deficiency

**DOI:** 10.1101/775775

**Authors:** Ajeet Chaudhary, Xia Chen, Jin Gao, Barbara Leśniewska, Richard Hammerl, Corinna Dawid, Kay Schneitz

## Abstract

Plant cells are encased in a semi-rigid cell wall of complex build. As a consequence, cell wall remodeling is essential for the control of growth and development as well as the regulation of abiotic and biotic stress responses. Plant cells actively sense physico-chemical changes in the cell wall and initiate corresponding cellular responses. However, the underlying cell wall monitoring mechanisms remain poorly understood. In Arabidopsis the atypical receptor kinase STRUBBELIG (SUB) mediates tissue morphogenesis. Here, we show that *SUB*-mediated signal transduction also regulates the cellular response to a reduction in the biosynthesis of cellulose, a central carbohydrate component of the cell wall. *SUB* signaling affects early increase of intracellular reactive oxygen species, stress gene induction as well as ectopic lignin and callose accumulation upon exogenous application of the cellulose biosynthesis inhibitor isoxaben. Moreover, our data reveal that *SUB* signaling is required for maintaining cell size and shape of root epidermal cells and the recovery of root growth after transient exposure to isoxaben. *SUB* is also required for root growth arrest in mutants with defective cellulose biosynthesis. Genetic data further indicate that *SUB* controls the isoxaben-induced cell wall stress response independently from other known receptor kinase genes mediating this response, such as *THESEUS1* or *MIK2*. We propose that *SUB* functions in a least two distinct biological processes: the control of tissue morphogenesis and the response to cell wall damage. Taken together, our results reveal a novel signal transduction pathway that contributes to the molecular framework underlying cell wall integrity signaling.

**Author Summary:** Plant cells are encapsulated by a semi-rigid and biochemically complex cell wall. This particular feature has consequences for multiple biologically important processes, such as cell and organ growth or various stress responses. For a plant cell to grow the cell wall has to be modified to allow cell expansion, which is driven by outward-directed turgor pressure generated inside the cell. In return, changes in cell wall architecture need to be monitored by individual cells, and to be coordinated across cells in a growing tissue, for an organ to attain its regular size and shape. Cell wall surveillance also comes also into play in the reaction against certain stresses, including for example infection by plant pathogens, many of which break through the cell wall during infection, thereby generating wall-derived factors that can induce defense responses. There is only limited knowledge regarding the molecular system that monitors the composition and status of the cell wall. Here we provide further insight into the mechanism. We show that the cell surface receptor STRUBBELIG, previously known to control organ development in Arabidopsis, also promotes the cell’s response to reduced amounts of cellulose, a main component of the cell wall.

## Introduction

Cell-cell communication is essential to regulate cellular behavior during many processes, including growth, development, and stress responses. In plants, the extra-cellular cell wall constitutes a central element of the underlying molecular mechanisms. It is mainly composed of carbohydrates, such as cellulose, hemicellulose, and pectin, and phenolic compounds, including lignin. Moreover, the cell wall also contains a plethora of different cell-wall-bound proteins [1,2]. It imposes restrictions on cell expansion and the movement of cells and serves as a barrier to pathogen attack. The cell wall counteracts turgor-driven growth and thus cell wall remodeling is required for cell expansion [3]. Cell wall fragments released by pathogen-derived lytic enzymes can act as danger signals and elicit plant immunity responses [4]. These observations imply a necessity for plant cells to monitor cell wall integrity (CWI). Such a mechanism would sense any physico-chemical alterations that occurred in the cell wall, and elicit a corresponding compensatory and protective cellular response [5–7].

Little is known about the molecular mechanisms that reside at the nexus of monitoring the cell wall status and the control of development and stress responses. Only a few cell surface signaling factors are presently implicated in monitoring CWI [5–7]. For example, a complex between RECEPTOR-LIKE PROTEIN44 (RLP44) and BRASSINOSTEROID INSENSITIVE1 (BRI1), the brassinosteroid receptor, specifically connects BRI1-mediated signaling to the detection of pectin modifications [8,9]. The extracellular domain of FERONIA (FER), a member of the *Catharanthus roseus* Receptor-like Kinase1-like (CrRLK1L) family of receptor kinases (RKs) originally identified on the basis of its role in sexual reproduction [10], binds pectin in vitro. *FER* is also required to prevent cell bursting upon exposure of root cells to salt [11]. Signaling mediated by ANXUR1 (ANX1) and ANX2, two other members of the CrRLK1L family, appears to contribute to monitoring cell wall integrity and the prevention of the premature burst of pollen tubes [12–15].

Plant cells also respond to changes in cellulose levels in the cell wall. Cellulose is present in the form of microfibrils that constitute the main load-bearing elements resisting turgor pressure. The microfibrils are embedded in matrix polysaccharides, mainly various hemicelluloses and pectins [1,2]. Cellulose is synthesized by cellulose synthase (CESA) complexes at the plasma membrane (PM) [16]. The effects of a reduced production of cellulose on plant growth and development can be studied by analyzing mutants with defects in genes encoding CESA subunits involved in primary cell wall biosynthesis [17–21]. Alternatively, pharmacological approaches can be applied. The herbicide isoxaben is a well-characterized inhibitor of cellulose biosynthesis [22,23]. A number of findings suggest CESAs to be the direct targets of isoxaben. First, several known isoxaben-resistant mutants carry mutations near the carboxyl terminus of certain CESA subunits [24,25]. Second, isoxaben induces a rapid clearing of CESA complexes from the PM [26]. Third, isoxaben uptake or detoxification appears unaffected in resistant plants [27].

The reaction of liquid culture-grown seedlings to isoxaben-induced cellulose biosynthesis inhibition (CBI) represents a thoroughly studied stress response to cell wall damage (CWD) [28–31]. The response is sensitive to osmotic support and eventually includes the upregulation of stress response genes, the production of reactive oxygen species (ROS), an accumulation of phytohormones, such as jasmonic acid (JA), changes in cell wall composition, including the production of ectopic lignin and callose, and finally growth arrest. Similar effects were also observed when studying the phenotypes of different *cesA* mutants [17–21].

The mechanism controlling the CWD response to CBI is known to involve three RKs [28]. THESEUS1 (THE1), another member of the CrRLK1L family, was first implicated in this process [32]. *THE1* was identified based on its genetic interaction with *PROCUSTE1* (*PRC1*), a gene encoding a CESA6 subunit [21]. Amongst others, cellulose-deficient *prc1* single mutants exhibit reduced hypocotyl length and ectopic lignin accumulation. In *the1 prc1* double mutants these effects are ameliorated although cellulose levels remain reduced [32]. Moreover, *THE1* is required for the altered expression levels of several stress-response genes upon exposing liquid-grown seedlings to isoxaben [33]. Recently, the leucine-rich repeat (LRR)-XIIb family RK MALE DISCOVERER1-INTERACTING RECEPTOR LIKE KINASE 2/LEUCINE-RICH REPEAT KINASE FAMILY PROTEIN INDUCED BY SALT STRESS (MIK2/LRR-KISS) [34,35] and the LRR-XIII family member FEI2 [36] have also been shown to participate in the isoxaben-induced cell wall stress response [28,33,37]. Genetic analysis revealed that *THE1* and *MIK2* have overlapping but also distinct functions, suggesting a complex regulation of the CBI response, with *THE1* and *MIK2* promoting this response via different mechanisms. *FEI2* appears to be part of the *THE1* genetic pathway [28].

Tissue morphogenesis in Arabidopsis requires signaling mediated by the atypical LRR-RK STRUBBELIG (SUB). SUB, also known as SCRAMBLED (SCM), is a member of the LRR-V family of RKs and controls several developmental processes, including floral morphogenesis, integument outgrowth, leaf development and root hair patterning [38–40]. SUB represents an atypical receptor kinase, as its *in vivo* function does not require enzymatic activity of its kinase domain [38,41]. Our previous studies indicate that SUB not only localizes to the PM but is also present at plasmodesmata (PD), channels interconnecting most plant cells [42,43], where it physically interacts with the PD-specific C2 domain protein QUIRKY (QKY) [44].

Current data also associate *SUB* signaling to cell wall biology. For example, whole-genome transcriptomics analysis revealed that many genes responsive to *SUB*-mediated signal transduction relate to cell wall remodeling [45]. Moreover, Fourier-transform infrared spectroscopy (FTIR)-analysis indicated that flowers of *sub* and *qky* mutants share overlapping defects in cell wall biochemistry [46]. Thus, apart from functionally connecting RK-mediated signal transduction and PD-dependent cell-cell communication *SUB* signaling also relates to cell wall biology.

Here, we report on a further exploration of the connection between the cell wall and SUB function. Our data reveal a novel role for *SUB* signaling in the CBI-induced CWD response. We show that *SUB* affects several processes, such as ROS accumulation, stress gene induction as well as ectopic lignin and callose accumulation, that are initiated upon application of exogenous isoxaben. Moreover, *SUB* signaling is necessary for maintaining cell shape and recovery of root growth after transient exposure to isoxaben. Our genetic data further indicate that *SUB, THE1*, and *MIK2* act in different pathways and that not all contributions of *SUB* to CBI-induced CWD signaling require *QKY* function.

## Results

In light of the connection between SUB signaling and cell wall biology, we set out to address if *SUB* plays a role in the seedling responses to cell wall stress. In particular, we focused on the possible role of *SUB* in the isoxaben-induced CWD response.

### *SUB* does not affect cellulose production

We first investigated if *SUB* influences cellulose biosynthesis in seven days-old seedlings. We first analyzed transcript levels of *CESA1, CESA3*, and *CESA6* in wild-type and *sub*. The three genes encode the CESA isoforms present in CSCs of the primary cell wall [47,48]. We could not detect differences in transcript levels of *CESA1, CESA3*, and *CESA6* between *sub* and wild-type in quantitative real-time polymerase chain reaction (qPCR) experiments (Fig. 1A). Moreover, we assessed the levels of cellulose and failed to detect differences between *sub* and wild-type (Fig. 1B). We could however detect a reduction in cellulose levels in *prc1-1* that was comparable to previous findings [21] (Fig. 1B). The results indicate that *SUB* does not play a central role in cellulose biosynthesis in seedlings.

**Fig 1.**
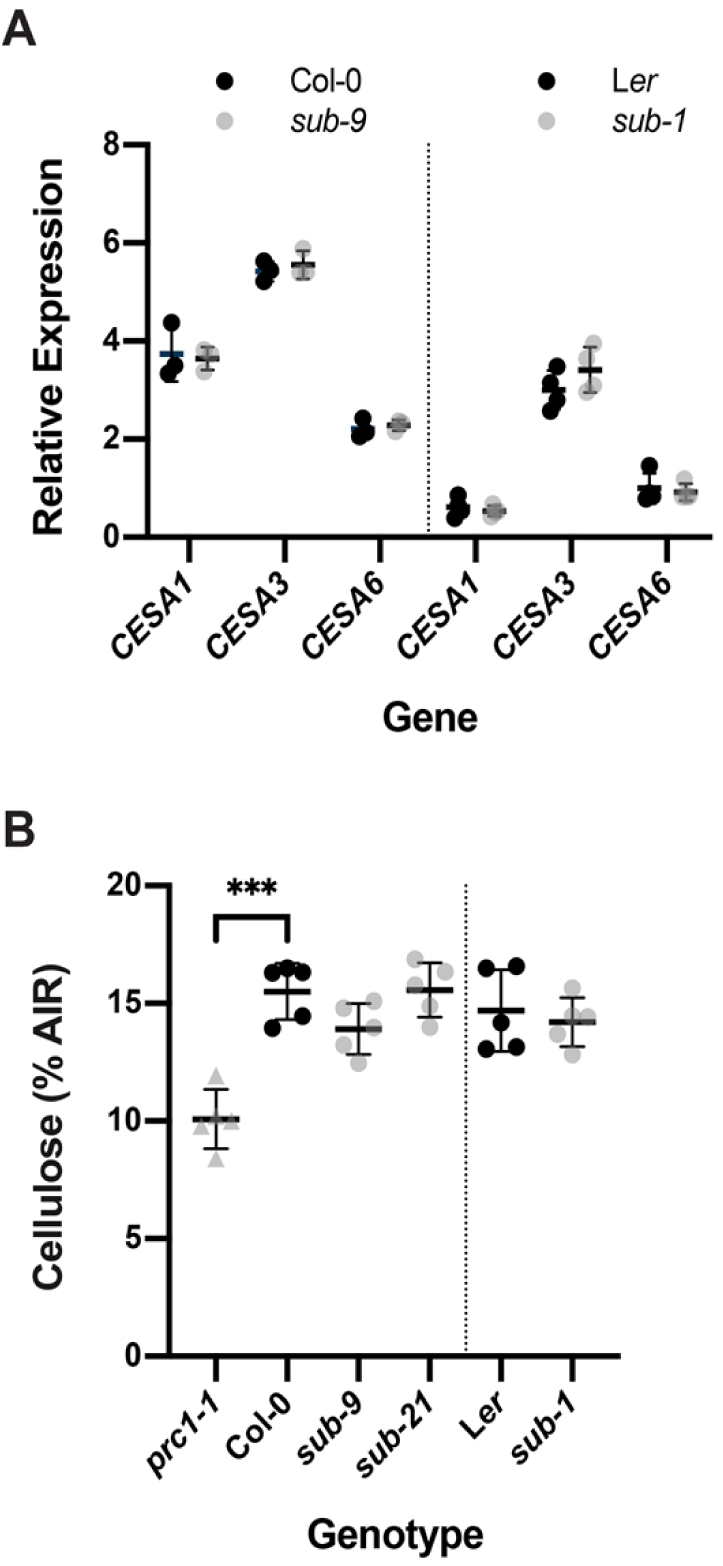
Effects of *SUB* on cellulose content in seven-day-old seedlings. (A) Gene expression levels for primary cell wall *CESA* genes in wild type and *sub* as assessed by qPCR. Individual *CESA* genes and genotypes are indicated. Mean ± SD is shown. Data points designate results of individual biological replicates. The experiment was performed two times with similar results. (B) Estimation of cellulose content. Genotypes are indicated. Data points indicate results of individual biological replicates. Mean ± SD is shown. Asterisks represent statistical significance (P < 0.0001, unpaired t test, two-tailed P values). No significant difference was observed between wild type and different *sub* mutants. Plants with a defect in *CESA6* (*prc1-1*) show a reduction in cellulose content as previously reported [21]. The experiment was repeated three times with similar results.

### *SUB* affects the isoxaben-induced CWD response

We then assessed if *SUB* activity is necessary for accumulation of reactive oxygen species (ROS) in response to isoxaben-induced CWD. To this end we exposed seven-day wild-type and *sub-9* seedlings, grown on half-strength Murashige and Skoog (MS) plates containing one percent sucrose, to 600 nM isoxaben in a time-course experiment. Seedlings were monitored for up to 120 minutes, at 30 minutes intervals. Upon treatment we assessed fluorescence intensity of the intracellular ROS probe H_2_DCFDA in roots [49,50]. In wild-type Col-0 seedlings treated for 30 minutes with isoxaben, we noticed an increase in H_2_DCFDA signal compared with untreated seedlings (Fig. 2A,B). Signal intensity of the probe increased further in seedlings exposed for 60 minutes. This signal intensity remained for up to 120 minutes of continuous exposure to isoxaben. In *sub-9* seedlings we detected a slightly increased H_2_DCFDA signal after 60 minutes (Fig. 2A,C). However, signal intensity was noticeably reduced in comparison to wild type. In comparison to the mock control, isoxaben-treated *sub-9* seedlings continued to show an enhanced signal intensity for up to 120 minutes of exposure, although the relative difference to signal levels of mock-treated seedlings was never as pronounced as in wild type. Thus in comparison to wild type, *sub-9* mutants showed a delayed onset of H_2_CDFDA signal appearance and an overall reduced signal intensity for the time frame analyzed. The results indicate that isoxaben causes the formation of intracellular ROS in roots of treated wild-type seedlings within 30 minutes of application. Moreover, *SUB* affects this ROS response.

**Fig. 2.**
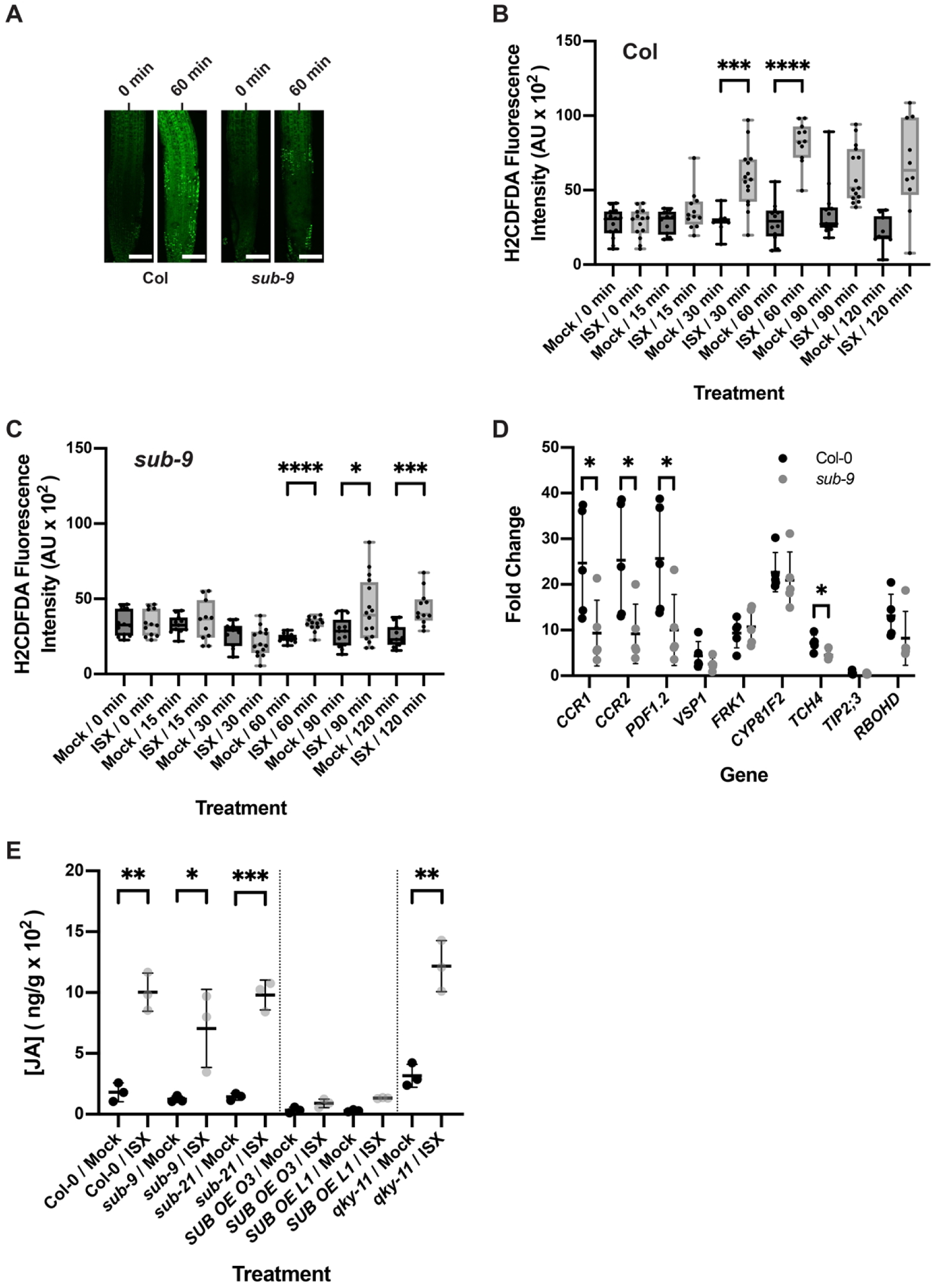
*SUB* effects on isoxaben-induced ROS production, marker gene expression and JA accumulation. (A) Confocal micrographs showing H_2_CFDA signal in root tips of six-day-old seedlings exposed to 600 nM isoxaben for the specified time. Genotypes are indicated. Note reduced signal in *sub-9*. (B, C) Quantification of results depicted in (A). Genotypes are indicated. Box and whisker plots are shown. 10 ≤ n ≤ 14. Asterisks represent statistical significance (_****_, P < 0.0001; _***_P < 0.002, _*_P < 0.05, unpaired t test, two-tailed P values). Experiments were performed three times with similar results. (D) Gene expression levels of several CBI marker genes by qPCR upon exposure of seven-day-old seedlings to 600 nM isoxaben for eight hours. The results from five biological replicates are shown. Marker genes and genotypes are indicated. Mean ± SD is presented. Asterisks represent statistical significance (_*,_ P < 0.05, unpaired t test, two-tailed P values). The experiment was repeated twice with similar results. (E) JA accumulation. Genotypes and treatments are indicated. Asterisks represent statistical significance (_***_, P < 0.002; _**_, P < 0.003; _*_P < 0.05, unpaired t test, two-tailed P values). n = 3. Experiments were repeated three times with similar results. Scale bars: (A) 100 μm.

Next, we tested if *SUB* activity is required for the transcriptional regulation of several marker genes, known to respond to isoxaben-induced CWD within eight hours [31]. We performed quantitative real-time polymerase chain reaction (qPCR) experiments using RNA isolated from seven days-old liquid-grown seedlings that had been incubated with 600 nM isoxaben for eight hours. We observed that isoxaben-induced upregulation of *CCR1, CCR2, PDF1.2*, and *TCH4* was attenuated in *sub-9* mutants compared to wild type (Fig. 2D). We did not detect a significant alteration in the upregulation of other tested marker genes, including *VSP1, FRK1, CYP81F2, TIP2;3*, and *RBOHD*, in *sub-9* seedlings. In wild type, upregulation of *CCR1* and *PDF1.2* expression was already detected upon four hours of isoxaben treatment [31]. We observed that expression levels of the two genes were significantly reduced in *sub-9* upon a four-hour exposure to isoxaben (Fig S1). These results indicate that *SUB* is required for the isoxaben-induced upregulation of expression of several marker genes.

The seedlings’ response to isoxaben also includes the accumulation of the phytohormone JA [31]. We thus tested if *SUB* affects the isoxaben-induced production of JA in seven days-old liquid-grown seedlings that had been incubated in 600 nM isoxaben for seven hours. We found that JA accumulation appeared largely unaffected in *sub-9* or *sub-21* mutants while the two overexpressing lines showed strongly diminished JA levels following isoxaben treatment (Fig. 2E). The results indicate that *SUB* is not required for isoxaben-induced JA accumulation. The observation that the phenotypes of the loss-of-function and overexpressing mutants are not easy to reconcile with each other renders an interpretation of the effects seen in the *SUB* overexpressing lines difficult. Thus, their biological relevance needs to be assessed in further experiments.

Isoxaben-induced CBI eventually results in the alteration of cell wall biochemistry as evidenced by the ectopic accumulation of lignin and callose [31]. To investigate if *SUB* affects lignin biosynthesis, we estimated lignin accumulation in roots using phloroglucinol staining after exposing six-day-old liquid-grown seedlings to 600 nM isoxaben for 12 hours. We observed reduced phloroglucinol staining in the root elongation zone of *sub-1* seedlings in comparison to wild type L*er* indicating less ectopic lignin production (Fig. 3A,B). We also noticed reduced phloroglucinol signal in *sub-9* seedlings (Col-0 background) although the effect was less prominent. However, in our hands Col-0 wild-type plants exhibited an overall weaker phloroglucinol staining indicating that isoxaben-induced lignin accumulation does not occur to the same level as in L*er* (Fig. 3A,B). We could detect increased phloroglucinol staining in two out of three *pUBQ::SUB:mCherry* lines (L1, O3) (Fig. 3A,B).

**Fig. 3.**
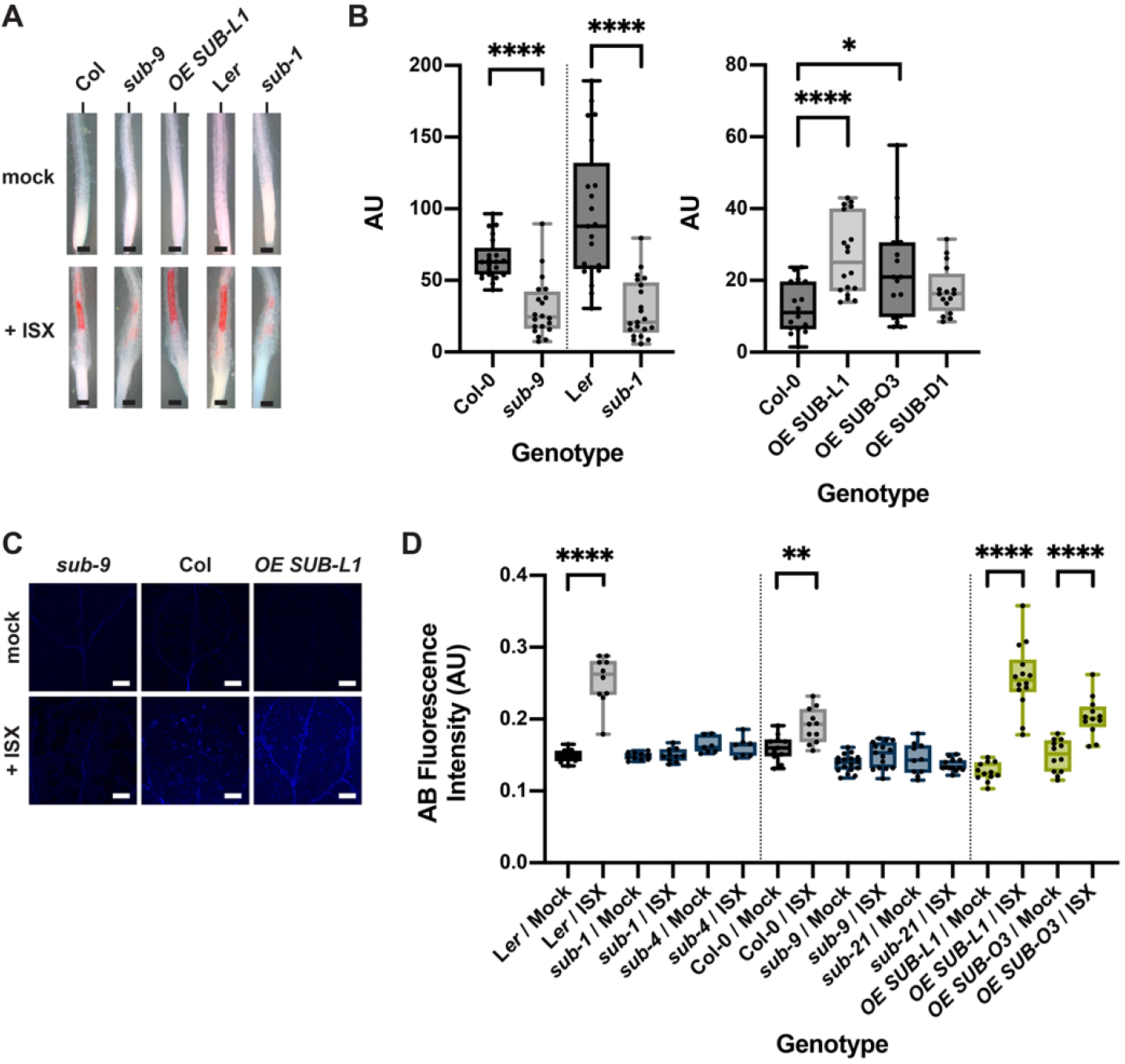
*SUB* affects isoxaben-induced lignin and callose accumulation. (A) Phloroglucinol signal strength indicating lignin accumulation in roots of six-day-old seedlings exposed to 600 nM isoxaben for 12 hours. Genotypes: Col, *sub-9* (Col), *pUBQ::gSUB:mCherry* (line L1), L*er*, and *sub-1* (L*er*). (B) Quantification of the results depicted in (A). Left panel shows results obtained from different *sub* mutants in the Col or L*er* background. Right panel depicts results from three independent *pUBQ::gSUB:mCherry* transgenic lines overexpressing *SUB* (Col, lines L1, O3, D1). Box and whisker plots are shown. 16 ≤ n ≤ 21. Asterisks represent statistical significance (_*_P < 0.02, _****_, P < 0.0001; unpaired t test, two-tailed P values). The experiment was performed three times with similar results. (C) Confocal micrographs show cotyledons of seven-day-old *sub-9*, Col, and *pUBQ::gSUB:mCherry* (line L1) seedlings treated with mock or 600 nM isoxaben for 24 hours. Aniline blue fluorescence signal strength indicates callose accumulation. (D) Quantification of the results depicted in (C). Left panel shows results obtained from *sub* mutants in L*er* background. Center panels indicates results obtained from *sub* mutants in Col background. Right panel depicts results from two independent *pUBQ::gSUB:mCherry* transgenic lines overexpressing *SUB* (Col, lines L1, O3). Box and whisker plots are shown. 7 ≤ n ≤ 18. Asterisks represent statistical significance (_****_, P < 0.0001; _**_P < 0.004, unpaired t test, two-tailed P values). The experiment was performed three times with similar results. Scale bars: (A) 0.1 mm; (C) 0.2 mm.

Isoxaben-treatment for 24 hours results in the formation of callose in cotyledons of wild-type seedlings [31]. Thus, we tested if *SUB* is required for this process as well. To this end we transferred seven days-old plate-grown seedlings to liquid medium without isoxaben for 12 hours. Subsequently, medium was exchanged, and seedlings were kept in 600 nM isoxaben for another 24 hours followed by callose detection using aniline blue staining [51]. As expected, we observed prominent aniline blue staining in cotyledons of L*er* and Col wild-type seedlings upon isoxaben treatment (Fig. 3C,D). By contrast, we detected strongly reduced aniline blue staining in cotyledons of isoxaben-treated *sub-1* and *sub-4* (both in L*er*) as well as *sub-9* and *sub-21* (both in Col). In contrast, and similar to the phloroglucinol staining described above, we detected stronger aniline blue staining in *SUB* overexpressors (lines L1, O3) (Fig. 3C,D).

Taken together, the results indicate that *SUB* is required for the isoxaben-induced formation of lignin and callose in seedlings.

### The *SUB*-mediated CBI response is sensitive to sorbitol

The isoxaben-induced CWD response is sensitive to turgor pressure, as indicated by the suppression of lignin or callose accumulation in the presence of osmotica, such as sorbitol [28,30,31]. To test if *SUB* affects a turgor-sensitive CBI response we compared isoxaben-induced accumulation of lignin and callose in six days-old Col-0, *sub-9*, and *pUBQ::SUB:mCherry* seedlings in co-treatments with 600 nM isoxaben and 300 mM sorbitol (Fig. 4A-C). We observed that sorbitol suppressed lignin and callose accumulation in all tested genotypes, including SUB:mCherry overexpressing lines, which show hyperaccumulation of lignin or callose in the absence of sorbitol.

**Fig. 4.**
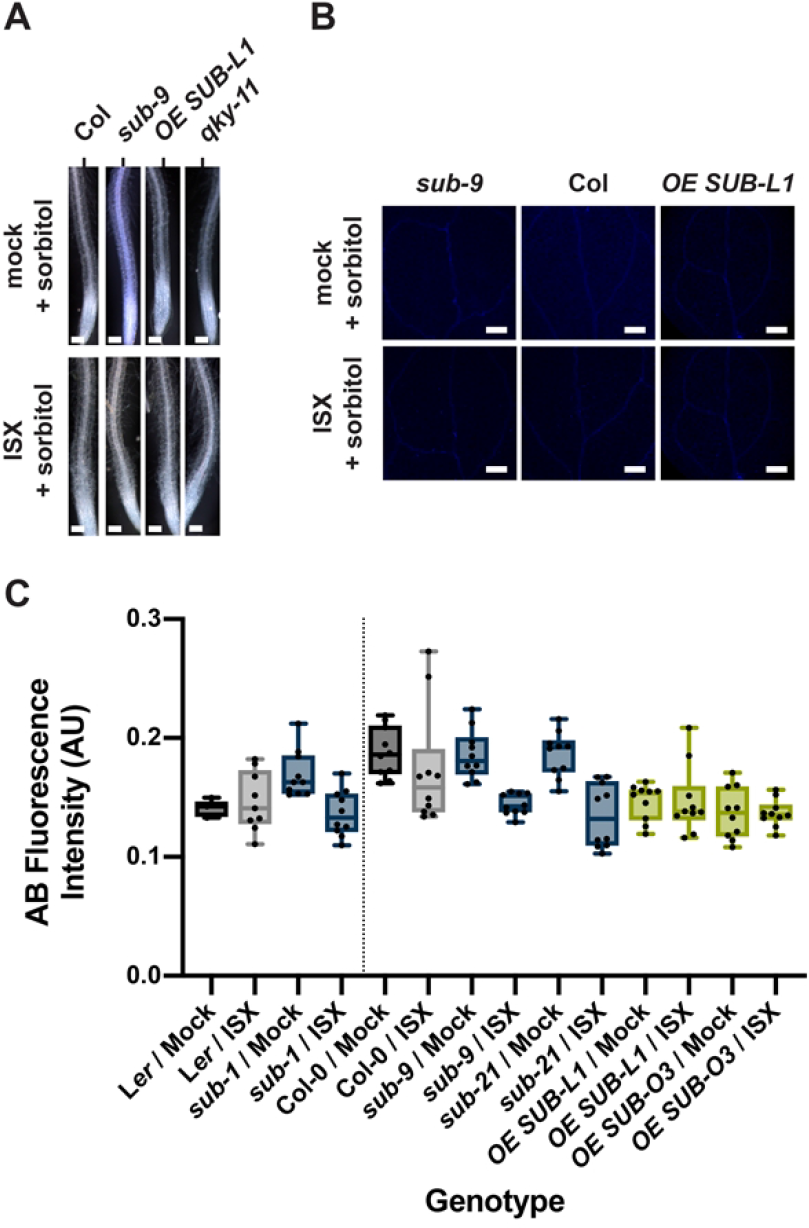
The effects of sorbitol on lignin and callose accumulation upon isoxaben exposure. (A) Phloroglucinol signal strength indicating lignin accumulation in roots of six-day-old seedlings exposed to mock/300 mM sorbitol or to 600 nM isoxaben/300 mM sorbitol for 12 hours. Genotypes: Col, *sub-9* (Col), *pUBQ::gSUB:mCherry* (line L1), *qky-11* (Col). Note absence of detectable signal. The experiment was performed three times with similar results (n ≥ 10). (B) Confocal micrographs show cotyledons of seven-day-old *sub-9*, Col, and *pUBQ::gSUB:mCherry* (Col, line L1) seedlings treated with mock/300 mM sorbitol or 600 nM isoxaben/300 mM sorbitol for 24 hours. Aniline blue fluorescence signal strength indicates callose accumulation. No increase in signal intensity can be observed in isoxaben-treated seedlings. (C) Quantification of the results depicted in (B). Left panel depicts results obtained from *sub-1* mutants in L*er* background. Right panel shows results obtained from *sub* mutants in Col background and also depicts results from two independent *pUBQ::gSUB:mCherry* transgenic lines overexpressing *SUB* (Col, lines L1, O3). Box and whisker plots are shown. 5 ≤ n ≤ 10. No statistically significant differences were observed (unpaired t tests, two-tailed P values). The experiment was performed three times with similar results. Scale bars: (A) 0.1 mm; (C) 0.2 mm.

### *SUB* attenuates isoxaben-induced cell swelling and facilitates root growth recovery

Next, we assessed the biological relevance of *SUB* in the isoxaben-induced CWD response. Exposure of seedlings to isoxaben eventually results in the shortening and swelling of cells of the root epidermis, possibly a result of reduced microfibril formation in the cell wall [28]. We transferred six days-old plate-grown seedlings into a mock solution or a solution containing 600 nM isoxaben for up to seven hours. We then assessed the timing of the initial appearance of altered cellular morphology of root epidermal cells. In addition, we monitored the severity of the phenotype. We focused on cells of the elongation zone that bordered the root meristem. Notably, we did not observe any obvious morphological alterations in mock-treated wild-type or mutant seedlings (Fig. 5). In Col-0 wild-type seedlings cell shortening and swelling first became noticeable during the five to six-hour interval proceeding treatment (Fig. 5) (44/96 seedlings total, n = 4), as reported previously [28]. Upon isoxaben application to *sub-9* or *sub-21* seedlings, however, similar cellular alterations were already detected at the three to four-hour interval post treatment initiation (*sub-9*: 23/57, n = 2; *sub-21*: 20/46, n = 2). In addition, *sub* mutants exhibited more pronounced cellular alterations after seven hours of isoxaben treatment in comparison to wild type (Fig. 5).

**Fig. 5.**
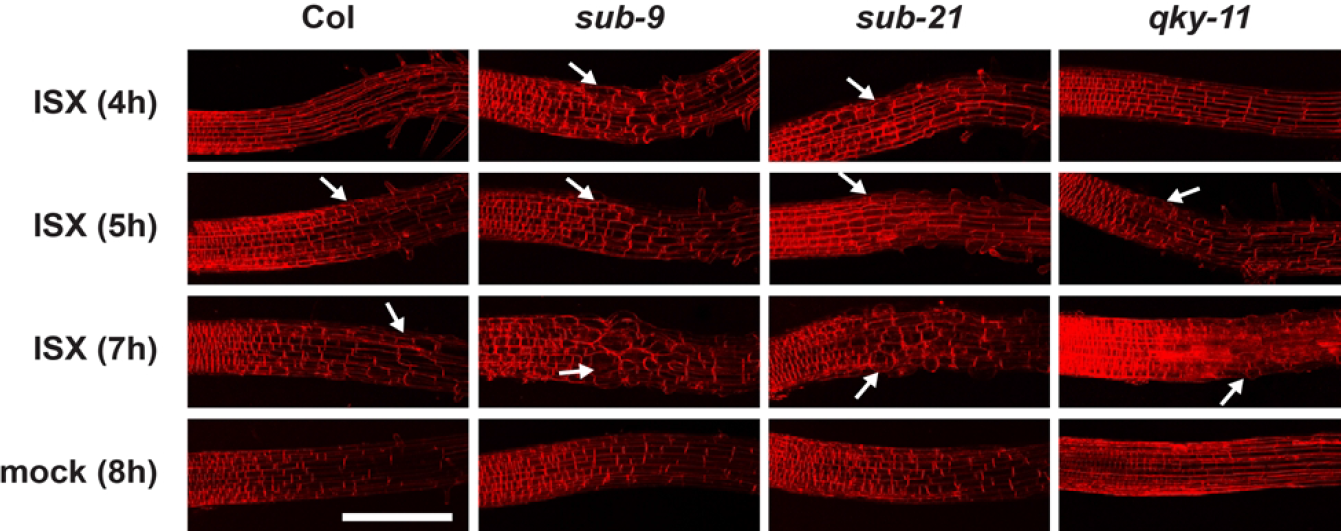
Root epidermal cell shape changes upon isoxaben treatment. Six-day-old seedlings counter-stained with the membrane stain FM4-64 are shown. Confocal micrographs depict the region where the elongation zone flanks the root meristem. Time of exposure in hours to 600 nM isoxaben (ISX) or mock is indicated as are the genotypes. Arrows denote aberrant cell shapes. Scale bar: 0.1 mm.

In wild-type seedlings, a 24-hour exposure to isoxaben results in a temporary stop of root growth followed by a rapid recovery [28]. We tested the role of *SUB* in root growth recovery upon a 24 hour-treatment with isoxaben. Six-day-old wild-type and mutant seedlings grown on plates were transferred onto media containing 600 nM isoxaben for 24 hours, then moved to fresh plates lacking isoxaben. Seedlings were then monitored for continued root growth at 24 hour intervals, for a total of 72 hours (Table 1). As control, we used the *ixr2-1* mutant, which is resistant to isoxaben due to a mutation in the *CESA6* gene [24,25,52]. We observed that 98 percent of *ixr2-1* seedlings recovered root growth already within 24 hours, indicating that treatment did not generally impact the seedlings’ ability to recover root growth. We then tested wild-type seedlings. We noticed that 46 percent of L*er* and 39 percent of Col seedlings had resumed root growth after 24 hours. By 72 hours, 86 percent of L*er* and 90 percent of Col seedlings had recovered root growth. In contrast, a significantly reduced fraction of *sub-1* and *sub-4* mutants had resumed root growth when compared to wild-type L*er* (Table 1). The *sub-9* and *sub-21* mutants also showed reduced root growth recovery in comparison to Col although *sub-21* appeared less affected than *sub-9*. Importantly, *ixr2-1 sub-9* mutants behaved identical to *ixr2-1* single mutants at all time points scored. These findings indicate that *ixr2-1* is epistatic to *sub-9* and that the observed isoxaben-induced decrease in root growth recovery in *sub-9* mutants relates to the herbicide.

**Table 1.** Root growth recovery after isoxaben treatment.

| Genotype | N total<sup>a</sup> | 24 h<sup>b</sup> | 48 h<sup>b</sup> | 72 h<sup>b</sup> |
| --- | --- | --- | --- | --- |
| *Ler* | 110 | 45.5 | 69.1 | 85.5 |
| *sub-1* | 82 | 24.4** | 50.0* | 64.6** |
| *sub-4* | 76 | 17.1** | 39.5**** | 52.6**** |
| *qky-8* | 87 | 37.9<sup>ns</sup> | 64.4<sup>ns</sup> | 88.5<sup>ns</sup> |
| Col-0 | 142 | 38.7 | 65.5 | 90.1 |
| *sub-9* | 88 | 23.9* | 45.5** | 70.5** |
| *sub-21* | 72 | 26.4<sup>ns</sup> | 51.4<sup>ns</sup> | 68.1** |
| *qky-11* | 81 | 37.0<sup>ns</sup> | 64.2<sup>ns</sup> | 92.6<sup>ns</sup> |
| *ixr2-1* | 83 | 97.6**** | 97.6**** | 97.6<sup>ns</sup> |
| *sub-9 ixr2-1* | 77 | 100.0**** | 100.0**** | 100.0** |
| *the1-1* | 78 | 41.0<sup>ns</sup> | 69.2<sup>ns</sup> | 88.5<sup>ns</sup> |

Percentages of root growth recovery of plate-grown, seven-day-old seedlings after transient exposure to 600 nM isoxaben for 24 hours. The top four sample rows list genotypes that are in *Ler* background. The bottom seven sample rows list genotypes that are in *Col* background.
<sup>a</sup>Total number of samples. Cases per class and timepoint were pooled from three independent experiments. For each biological replicate $22 \leq n \leq 32$ seedlings were analyzed per genotype and time point.
<sup>b</sup>Statistical significance (mutant vs respective wild type): na, not applicable; ns, not significant; \* $P < 0.05$ ; \*\* $P < 0.005$ ; \*\*\*\* $P < 0.0001$ ; (Fisher's exact test; two-sided).

Taken together, the results suggest that *sub* mutants are hypersensitive to isoxaben treatment and that *SUB* facilitates root growth recovery, and represses cell size and shape changes in root epidermal cells during the isoxaben-induced CWD response.

### *SUB* attenuates root growth in *prc1-1*

*PRC1* encodes the CESA6 subunit of cellulose synthase [21] and *prc1* loss-of-function mutants show reduced cellulose levels [21] (Fig. 1B). In addition, *prc1-1* mutants are characterized by a reduced elongation of etiolated hypocotyls and roots [21]. To test if *SUB* also affects a biological process in a scenario where cellulose reduction is induced genetically we compared root length in *sub-9, sub-21*, and *prc1-1* single and *sub-9 prc1-1* double mutants (Fig. 6A,B). We found that root length of *sub-9* or *sub-21* did not deviate from wild type while root length of *prc1-1* was markedly smaller in comparison to wild type, confirming previous results [21]. Interestingly, however, we observed that *sub-9 prc1-1* exhibited a significantly longer root than *prc1-1* though *sub-9 prc1-1* roots were still notably smaller than wild type roots. The results indicate that *SUB* contributes to root growth inhibition in *prc1-1*.

**Fig. 6.**
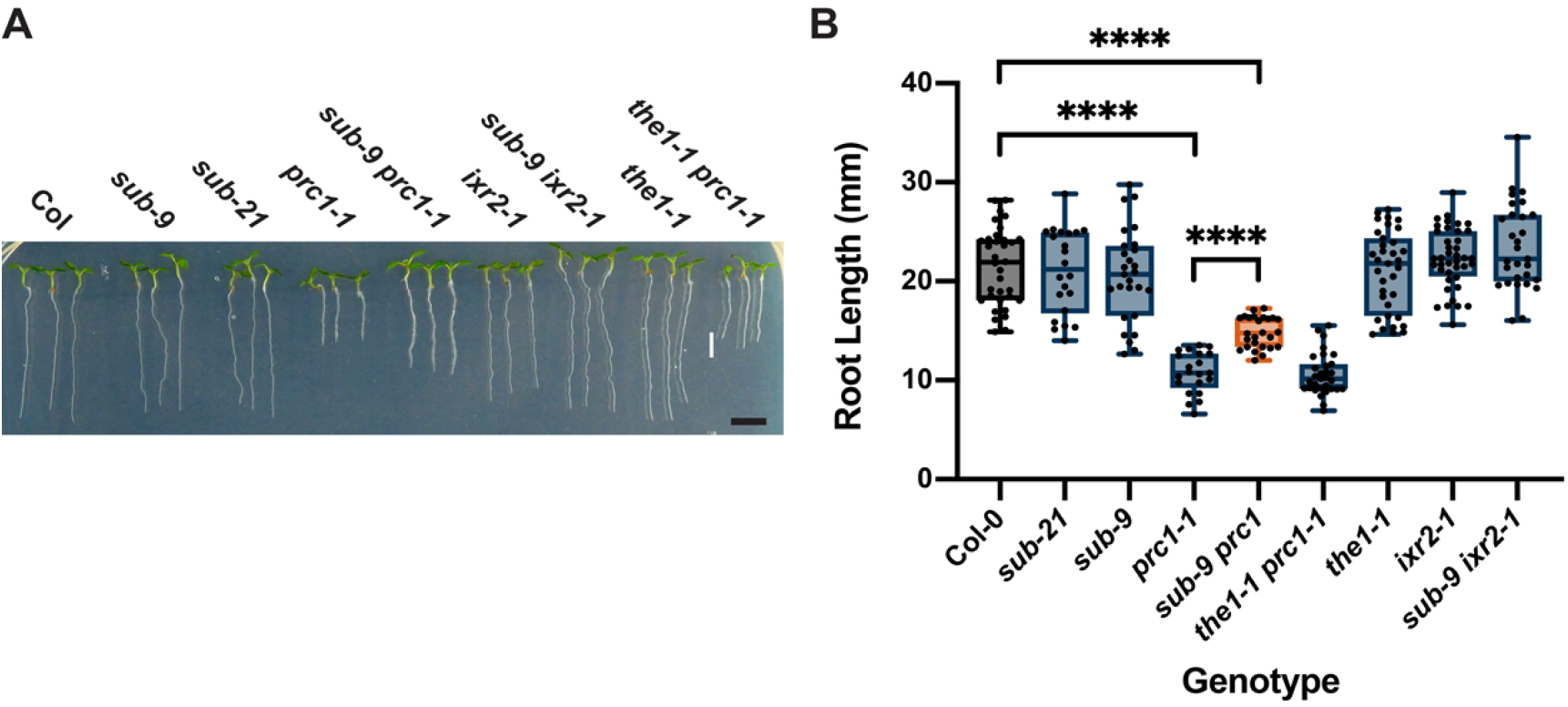
*SUB* effect on root growth inhibition in *prc1-1*. (A) Root length in seven-day-old seedlings grown on plates under long-day conditions (16 hours light). Genotypes are indicated. Note the partial rescue of root length in *sub-9 prc1-1* but not in *the1-1 prc1-1*. (B) Quantification of the data shown in (A). Box and whisker plots are shown. 21 ≤ n ≤ 40. Asterisks represent statistical significance (_****_, P < 0.0001; unpaired t test, two-tailed P values). The experiment was performed three times with similar results. Scale bars: 0.5 mm.

### *SUB* and *QKY* contribute differently to the CBI response

Evidence suggests that a protein complex including SUB and QKY is important for SUB-mediated signal transduction regulating tissue morphogenesis [44,53,54]. Thus, we wanted to explore if *QKY* is also required for the isoxaben-induced CWD response in seedlings. We first investigated if *QKY* affects the early isoxaben-induced changes in intracellular ROS levels by assessing H_2_CDFDA fluorescence in root tips of *qky-11* seedlings that were treated with 600 nM isoxaben. We did not observe a difference in signal intensity between mock and isoxaben-treated *qky-11* (Fig. 7A). Moreover, signal intensity was similar to wild type (Fig. 2B) indicating that *QKY* does not contribute to altered intracellular ROS levels in root tips of treated seedlings in a noticeable fashion. We then tested if *QKY* promotes isoxaben-induced marker gene expression in liquid-grown seedlings. Using qPCR we observed that *SUB* and *QKY* affect a similar set of marker genes (Fig. 7B). Next, we assessed isoxaben-induced lignin accumulation in wild-type and *qky* seedlings by phloroglucinol staining. We noticed reduced staining in *qky-8* and *qky-11* mutants compared to wild type (Fig. 7C,D). Again, the effect was less obvious in Col-0. In addition, we noticed that *qky-11* did not affect the absence of lignin accumulation in seedlings simultaneously treated with isoxaben and sorbitol (Fig. 4A). We also investigated the role of *QKY* in isoxaben-induced callose deposition by scoring the aniline blue-derived signal in cotyledons. In contrast to *sub* mutants, however, we did not observe a significant difference in signal strength between *qky-8, qky-11*, and wild type (Fig. 7E,F). Regarding isoxaben-induced JA accumulation we found that *qky-11* seedlings did not noticeable deviate from *sub* (Fig. 2E). We also analyzed isoxaben-induced shortening and swelling of root epidermal cells in *qky-11* mutants (Fig. 4). We noticed the first defects in a four to five hour interval and thus about an hour later than in *sub* mutants (37/63, n = 3). Cell swelling after seven hours exposure to isoxaben was prominent in *qky-11* but less severe in comparison to *sub-9* (Fig. 4). Finally, we investigated root growth recovery after transient isoxaben application. We observed that *qky-8* and *qky-11* mutants did not significantly deviate from wild type (Table 1).

**Fig. 7.**
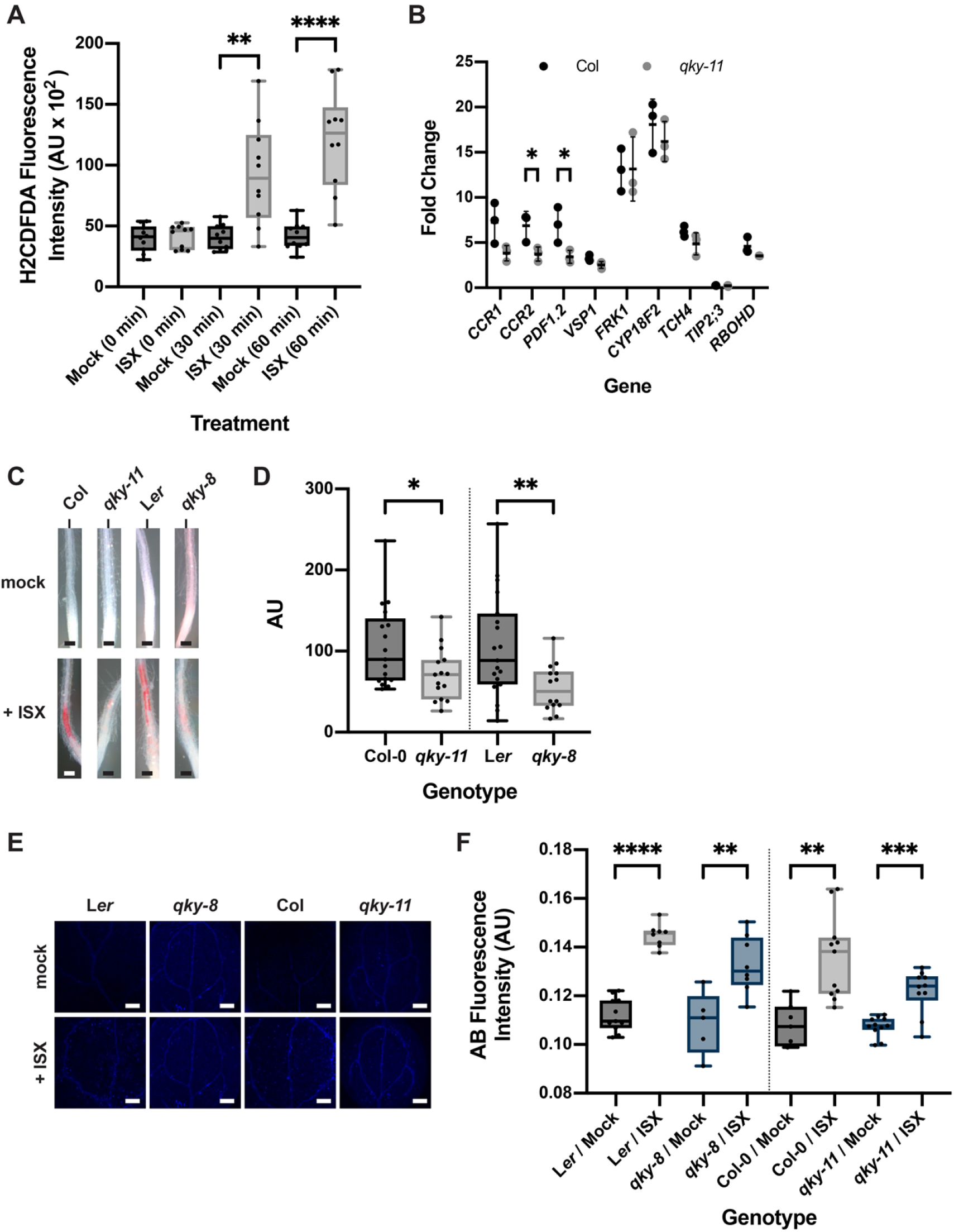
Role of *QKY* in isoxaben-induced CBI responses. (A) Quantification of H_2_CDFDA signal indicating ROS accumulation in root tips of six-day-old *qky-11* seedlings exposed to mock or 600 nM isoxaben for the indicated time. Note the unaltered signal. Box and whisker plots are shown. n = 10. Asterisks represent statistical significance (_**,_ P = 0.0013; _****_, P < 0.0001, unpaired t test, two-tailed P values). Experiments were performed three times with similar results. (B) Gene expression levels of several CBI marker genes by qPCR upon exposure of seven-day-old seedlings to 600 nM isoxaben for eight hours. The results from three biological replicates are shown. Marker genes and genotypes are indicated. Mean ± SD is presented. Asterisks represent statistical significance (_*,_ P < 0.05, unpaired t test, two-tailed P values). The experiment was repeated three times with similar results. (C) Phloroglucinol signal strength indicating lignin accumulation in roots of six-day-old seedlings exposed to 600 nM isoxaben for 12 hours. Genotypes: Col, *qky-11* (Col), L*er*, and *sub-1* (L*er*). (D) Quantification of the results depicted in (C). Genotypes are indicated. Box and whisker plots are shown. 15 ≤ n ≤ 19. Asterisks represent statistical significance (_*_P < 0.04, _**_, P < 0.01; unpaired t test, two-tailed P values). The experiment was performed three times with similar results. (E) Confocal micrographs show cotyledons of seven-day-old L*er, qky-8* (L*er*), Col, and *qky-11* (Col) seedlings treated with mock or 600 nM isoxaben for 24 hours. Aniline blue fluorescence signal strength indicates callose accumulation. (F) Quantification of the results depicted in (E). Left panel shows results obtained from *qky-8* mutants in L*er* background. Right panel indicates results obtained from *qky-11* mutants in Col background. Box and whisker plots are shown. 5 ≤ n ≤ 11. Asterisks represent statistical significance (_**_P < 0.006, ***, P = 0.0001; _****_, P < 0.0001; unpaired t test, two-tailed P values). The experiment was performed three times with similar results. Scale bars: (C) 0.1 mm; (E) 0.2 mm.

Taken together, the results indicate that *QKY* and *SUB* contribute to the isoxaben-mediated induction of an overlapping set of marker genes as well as to lignin accumulation. Moreover, *QKY* also plays a role in the suppression of isoxaben-induced alterations in cell morphology in the root epidermis. However, the results also imply that *QKY* is not required for isoxaben-induced early ROS accumulation in root tips as well as callose accumulation in cotyledons and does not affect root growth recovery after transient exposure to isoxaben.

### *SUB* and *THE1* share partially overlapping functions

*THE1* is a central regulator of the isoxaben-induced CBI response [28,33,37]. Our data indicate that *SUB* and *THE1* have overlapping but also distinct functions in this process. For example, *SUB* and *THE1* control isoxaben-induced lignin accumulation in roots (Fig. 3) [28,33]. However, in contrast to *THE1, SUB* was not shown to affect *FRK1* induction by isoxaben (Fig. 2D) [33]. We also found that root growth recovery of *the1-1* seedlings upon transient exposure to isoxaben did not deviate from wild type (Table 1). Moreover, we failed to detect an effect of *THE1* on root growth inhibition in *prc1-1*, while *SUB* contributes to this process (Fig. 6). We therefore explored further the relationship between *THE1* and *SUB*.

*THE1* contributes to the reduction of hypocotyl length of cellulose-diminished *prc1-1* seedlings grown in the dark as evidenced by the partial recovery of hypocotyl elongation in *the1 prc1* double mutants [32]. We compared *sub-9 prc1-1* to *the1-1 prc1-1* with respect to hypocotyl elongation in six-day-old etiolated seedlings and root length in seven-day-old light-grown seedlings (Fig. 8). We observed strongly reduced hypocotyl elongation in *prc1-1* in comparison to wild type and a significant suppression of this reduction in *the1-1 prc1-1* double mutants (Fig. 8A,B), as described earlier [32]. In contrast, hypocotyl length in *sub-9* mutants was not decreased, nor was there a partial reversal of reduced hypocotyl elongation in *sub-9 prc1-1* double mutants (Fig. 8A,B). The results indicate that *SUB* does not affect hypocotyl elongation in etiolated wild-type or *prc1-1* seedlings.

**Fig. 8.**
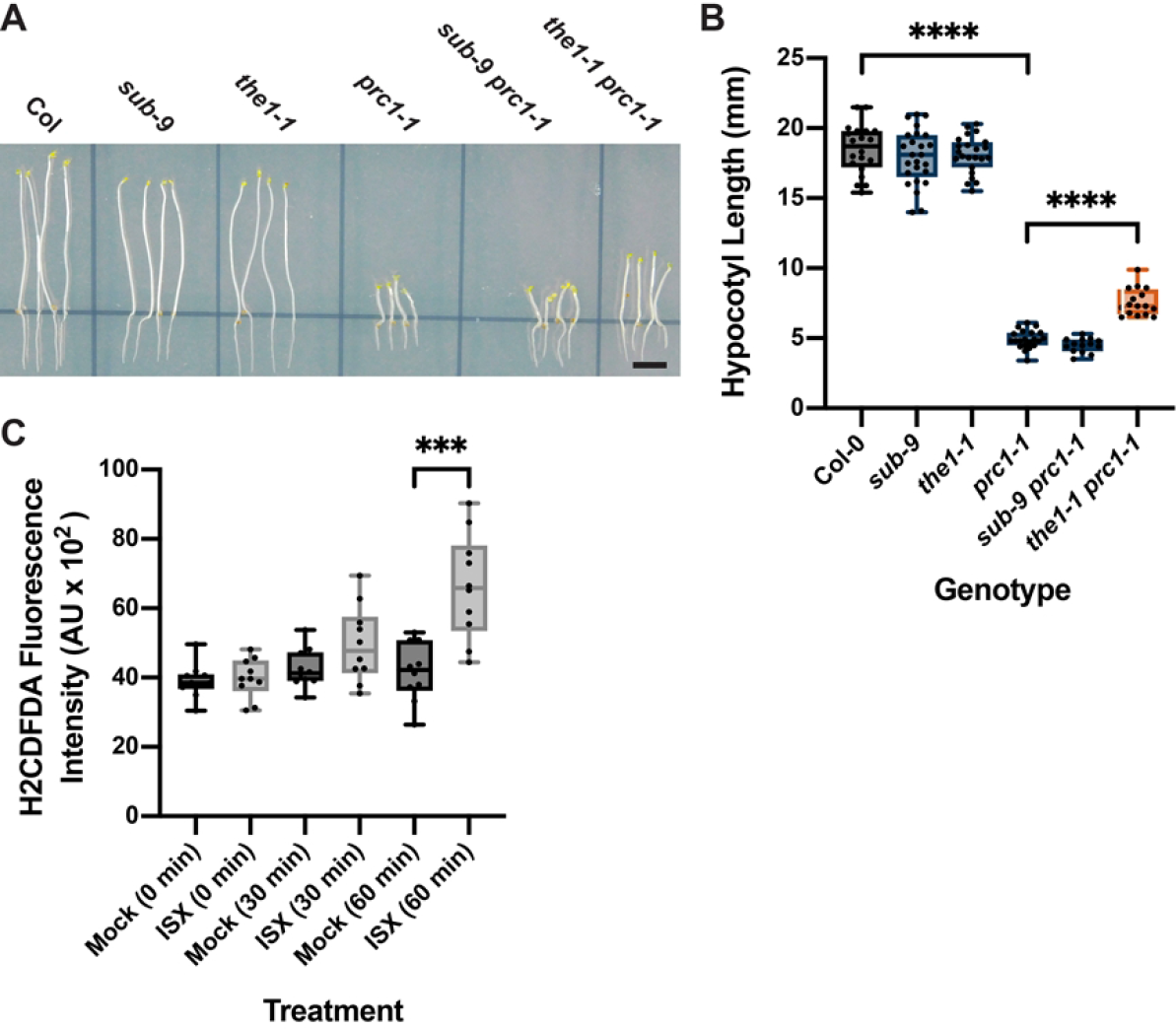
Influence of *THE1* on etiolated hypocotyl length of *prc1-1* and isoxaben-induced ROS accumulation in root tips. A) Hypocotyl elongation in six-day-old seedlings grown on plates in the dark. Genotypes are indicated. Note the partial rescue of hypocotyl elongation in *the1-1 prc1-1* but not in *sub-9 prc1-1*. (B) Quantification of the data shown in (A). Box and whisker plots are shown. 14 ≤ n ≤ 25. Asterisks represent statistical significance (_****_, P < 0.0001; unpaired t test, two-tailed P values). The experiment was performed three times with similar results. (C) Quantification of H_2_CDFDA signal indicating ROS accumulation in root tips of six-day-old *the1-1* seedlings exposed to mock or 600 nM isoxaben for the indicated time. Note the unaltered signal at 30 minutes exposure. Box and whisker plots are shown. n = 10. Asterisks represent statistical significance (_***_, P = 0.0003, unpaired t test, two-tailed P values). Experiments were performed three times with similar results. Scale bars: 0.5 mm.

Finally, we assessed the role of *THE1* in isoxaben-induced early ROS accumulation in root tips. We noticed no difference in H_2_CDFDA signal intensity when comparing *the1-1* seedlings that had been treated with either mock or isoxaben for 30 minutes (Fig. 8C). By contrast, isoxaben-induced reporter signal in the root tip became stronger than the mock-mediated signal at the 60 minutes time point (Fig. 8C). Overall the results resemble the findings obtained with *sub-9* mutants (Fig. 2C) although the response may be slightly less affected in *the1-1* in comparison to *sub-9*.

In summary, *SUB* and *THE1* are required for ROS and lignin accumulation in roots and show partial overlap with respect to stress marker gene induction. *SUB* and *THE1* show opposite effects on growth inhibition of roots and hypocotyls in *prc1* mutants and their functions differ with respect to root growth recovery.

## Discussion

Cell wall signaling during plant development and stress responses relies on complex and largely unknown signaling circuitry [5–7,29,55]. Only a few RKs, including THE1, MIK2, and FEI2, have so far been shown to play a major role in CBI-induced CWD signaling [28,32,33,37]. Our data establish *SUB* signal transduction as a novel component in the molecular framework mediating the CWD response.

To date, published evidence has attributed a developmental role to *SUB*, particularly in the control of tissue morphogenesis and root hair pattern formation [38–40,45,56]. The evidence provided in this work identifies a novel role for *SUB* in CWD signaling. Application of isoxaben results in reduced levels of cellulose [22,23]. Our results indicate that *SUB* affects multiple aspects of the isoxaben-induced CWD response. The observation that *sub-9* partially suppresses reduced root length exhibited by *prc1-1* indicates that *SUB* also mediates a CWD response that is caused by a genetic reduction in cellulose content. Thus, the collective data support the notion that the origin of the CWD relates to the reduced production of cellulose, a major carbohydrate component of the cell wall, and that *SUB* contributes to the compensatory cellular response to this type of cell-wall-related stress.

The data indicate that *SUB* already affects the early response to isoxaben-induced CWD, in particular ROS production. Previous results had revealed that the isoxaben-induced CWD response involves *THE1*-dependent ROS production [31,57]. In luminol-based extracellular ROS assays involving entire seedlings ROS production could be detected after around three to four hours following the application of isoxaben [57]. We used the intracellular ROS probe H_2_CDFDA and a microscope-based method that enabled tissue-level resolution. H_2_CDFDA has for example been used to assess basal intracellular ROS levels when studying *GLYCERALDEHYDE-3-PHOSPHATE DEHYDROGENASE* (*GAPDH*) genes and root hairs [49,50]. Our time-course data indicate that isoxaben induces the formation of intracellular ROS in the root meristem within 30 minutes. To our knowledge the isoxaben-dependent change in H_2_CDFDA fluorescence signal represents the earliest available marker for the isoxaben-induced CWD response. It also indicates that this response occurs much earlier than previously appreciated.

The results suggest that *SUB* is required for full induction of several marker genes, such as *CCR1, PDF1.2*, or *TCH4*. Moreover, the data indicate that *SUB* promotes CBI-induced accumulation of lignin and callose. In addition, *SUB* does not obviously influence the reduced lignin and callose state the results from double application of sorbitol and isoxaben. This result indicates that *SUB* mediates an osmosensitive CWD response. It has been proposed that a mechanical stimulus initiates the CWD response with the reduction in cellulose and corresponding weakening of the cellulose framework counteracting turgor pressure. In this model, a displacement or distortion of the plasma membrane relative to the cell wall could take place [28,58–61]. We speculate that *SUB* signaling likely involves as yet unknown mechano-sensitive factors. The combined results fulfil the criteria that have been established for a CBI-induced CWD response [28,31].

Interestingly, different loss-of-function (*sub-1, sub-9, sub-21*) or gain-of-function (*pUBQ::SUB:mCherry*) mutants show reciprocal effects regarding lignin and callose accumulation, with *sub* mutants showing less and several independent *pUBQ::SUB:mCherry* lines exhibiting higher levels of lignin or callose, respectively, upon application of isoxaben. Based on this evidence, we propose the model that *SUB* represents an important genetic regulator of isoxaben-induced lignin and callose accumulation and thus cell wall composition.

Several lines of evidence indicate that *SUB* plays a biologically relevant role in CWD signaling initiated by a reduction in cellulose content. Firstly, *SUB* attenuates isoxaben-induced cell bulging in the epidermal cells of the meristem-transition zone boundary of the root. Secondly, *SUB* facilitates root growth recovery upon transient exposure of seedlings to isoxaben. Thirdly, *SUB* is involved in root growth inhibition that is a consequence of a genetic reduction of cellulose content. In particular, root length of *sub-9 prc1-1* double mutant seedlings is less diminished in comparison to the root length of *prc1-1* single mutant seedlings. The results imply that *SUB* contributes to a compensatory response that counteracts the cellular and growth defects caused by reduced cellulose synthesis and further support the notion that *SUB* plays a central role in CBI-induced CWD signaling.

How does SUB relate to other known RK genes mediating the response to CBI-induced cell wall stress, such as *THE1* and *MIK2*? *SUB, THE1*, and *MIK2* all promote isoxaben-induced ectopic lignin production. The three genes are also required for full induction of certain stress marker genes. However, while *THE1* and *MIK2* control *FRK1* expression, and *MIK2* also affects the expression of *CYP81F2* [33], our data indicate that *SUB* is not required for the induction of these two genes. In addition, *THE1* and *MIK2* are necessary for isoxaben-induced JA accumulation [28,33,57], a process that apparently does not require *SUB* function. We also did not observe an effect of *THE1* on root growth recovery upon transient exposure to isoxaben. In addition, we did not find an altered hypocotyl length of *sub-9 prc1-1* in comparison to *the1-1* indicating that *SUB* does not affect hypocotyl growth inhibition in etiolated *prc1-1* seedlings, in contrast to *THE1* [32]. At the same time, our data suggest that *SUB* contributes to root growth inhibition in *prc1-1*. However, our evidence does not support a function for *THE1* in this process, as root length of *the1-1 prc1-1* double mutants did not deviate from the root length observed for *prc1-1* single mutants. This finding also implies that the lignin accumulation in the mature root parts of *prc1-1* (suppressed in *prc1-1 the1-1* double mutants) [32] does not correlate with root growth inhibition. Finally, we did not observe the left-hand root skewing in *sub* seedlings that is characteristic for *mik2* mutants [33], and we failed to observe floral defects in *mik2* or *the1* mutants. Taken together, the data suggest that *SUB* has both overlapping and distinct functions with *THE1* and *MIK2*. As the most parsimonious explanation of our results, we propose that *SUB* contributes to CBI-induced cell wall damage signaling independently from *THE1* and *MIK2* signaling, however the signaling pathways downstream of the different cell surface receptor kinases eventually partially converge and contribute to a subset of overlapping downstream responses.

*QKY* represents a central genetic component of *SUB*-mediated signal transduction during tissue morphogenesis and root hair patterning [45]. The expression pattern of *QKY* and *SUB* fully overlap [44] and present evidence supports the notion that SUB and QKY are part of a protein complex with QKY likely acting upstream to, or in parallel with, SUB [44,53,54]. However, genetic and whole-genome transcriptomic data suggested that *SUB* and *QKY* also play distinct roles during floral development [45]. The data presented in this study reveal that *SUB* and *QKY* both contribute to CBI-induced CWD signaling. Similar to *SUB QKY* is required for full induction of the same marker genes and for lignin accumulation. Moreover, *QKY* is also necessary for the prevention of cell bulging in the epidermal cells of the meristem-transition zone boundary of the root although the weaker phenotype of *qky-11* mutants indicates a lesser requirement for *QKY* in comparison to *SUB*. However, our data also indicate that *QKY* is not required for early ROS accumulation, ectopic callose accumulation, and root growth recovery as we found these three processes to be not noticeably affected in *qky* mutants. Thus, these data provide genetic evidence that the functions of *SUB* and *QKY* do not completely overlap and that *SUB* exerts CWD signaling functions that are independent of *QKY*. One way to rationalize these findings is to assume that SUB can also function in isolation or in protein complexes or pathways that do not involve QKY.

The diverse functions of *SUB* in development and the CWD response are likely to be achieved through participation in different signaling pathways. It is not uncommon that RKs play important roles in several biological processes. In this respect, SUB resembles for example the RK BRASSINOSTEROID INSENSITIVE 1-ASSOCIATED KINASE 1 (BAK1) / SOMATIC EMBROYGENESIS RECEPTOR KINASE 3 (SERK3), which functions in growth, development, and plant defense [62,63]. BAK1 interacts with a range of different LRR-RKs, including FLAGELLIN SENSING 2 (FL2) and BRASSINOSTEROID INSENSITIVE 1 (BRI1), and the discrimination between the growth and immunity functions of BAK1 was recently shown to rely on phosphorylation-dependent regulation [64,65]. In light of these considerations it is reasonable to propose that SUB is a member of different receptor complexes. As kinase activity of SUB is not required for its function [38] SUB could act as a scaffold around which the components of the various complexes assemble. A scaffold role has for example been proposed for the RK AtCERK1/OsCERK1 in chitin signaling or the RK FER in immune signaling [66,67]. It will be interesting to explore this notion in future work.

## Materials and Methods

### Plant work, plant genetics and plant transformation

*Arabidopsis thaliana* (L.) Heynh. var. Columbia (Col-0) and var. Landsberg (*erecta* mutant) (L*er*) were used as wild-type strains. Plants were grown as described earlier [45]. The *sub-1, qky-8* (all in Ler), and the *sub-9* and *qky-11* mutants (Col) have been characterized previously [38,41,44,45]. The *prc1-1* [21], *the1-1* [32], and *ixr2-1* [24] alleles were also described previously. The *sub-21* (Col) allele was generated using a CRISPR/Cas9 system in which the egg cell-specific promoter pEC1.2 controls Cas9 expression [68]. The single guide RNA (sgRNA) 5’-TAATAACTTGTATATCAACTT-3’ binds to the region +478 to +499 of the *SUB* coding sequence. The sgRNA was designed according to the guidelines outlined in [69]. The mutant carries a frameshift mutation at position 495 relative to the *SUB* start AUG, which was verified by sequencing. The resulting predicted short SUB protein comprises 67 amino acids. The first 39 amino acids correspond to SUB and include its predicted signal peptide of 29 residues, while amino acids 40 to 67 represent an aberrant amino acid sequence. The pUBQ::gSUB:mCherry plasmid used to generate the *SUB* overexpression lines L1 and O3 was generated previously [44]. Wild-type, and *sub* mutant plants were transformed with different constructs using Agrobacterium strain GV3101/pMP90 [70] and the floral dip method [71]. Transgenic T1 plants were selected on Kanamycin (50 μg/ml), Hygromycin (20 μg/ml) or Glufosinate (Basta) (10 μg/ml) plates and transferred to soil for further inspection.

### Cellulose quantification

Seedlings were grown on square plates with half strength MS medium and 0.3% sucrose supplemented with 0.9% agar for seven days. Cellulose content was measured following the Updegraff method essentially as described [72,73], with minor modifications as outlined here. Following the acetic nitric treatment described in [72], samples were allowed to cool at room temperature and transferred into 2 ml Eppendorf safety lock tubes. Samples were then centrifuged at 14000 rpm at 15°C for 15 min. The acetic nitric reagent was removed carefully without disturbing the pelleted material at the bottom of the tube. 1 ml of double-distilled H_2_O was added, and the sample was left on the bench for 10 min at room temperature followed by centrifugation at 14000 rpm at 15°C for 15 min. After aspirating off the H_2_O 1 ml acetone was added and the samples were incubated for another 15 min, followed by centrifugation at 14000 rpm at 15°C for 15 min. Afterwards acetone was removed, and samples were air-dried overnight. Then the protocol was continued as described in [72].

### PCR-based gene expression analysis

For quantitative real-time PCR (qPCR) of *CesA* and stress marker genes 35 to 40 seedlings per flask were grown in liquid culture under continuous light at 18°C for seven days followed by treatment with mock or 600 nM isoxaben for eight hours or on plates (21°C, long-day conditions). With minor changes, RNA extraction and quality control were performed as described previously [74]. cDNA synthesis, qPCR, and analysis were done essentially as described [75]. Primers are listed in Supplemental Table 1.

### ROS, lignin, and callose staining

Intracellular ROS accumulation in root meristems was estimated using the H_2_DCFDA fluorescent stain essentially as described [50]. Seeds were grown on square plates with half strength MS medium and 1% sucrose supplemented with 0.9% agar. The seeds were stratified for two days at 4°C and incubated for seven days at 22°C under long day conditions, at a 10 degree inclined position. Seven days-old seedlings were first transferred into multi-well plates containing half strength liquid MS medium supplemented with 1% sucrose for two hours. Then medium was exchanged with liquid medium containing either DMSO (mock) or 600nM isoxaben without disturbing seedlings. 10 min prior to each time point seedlings were put in the dark and the liquid medium was supplemented with 100 μM H_2_DCFDA staining solution. Images was acquired with a confocal microscope. For quantification a defined region of interest (ROI) located 500 μm above the root tip (excluding the root cap) was used in all samples. Staining for lignin (phloroglucinol) and callose (aniline blue) was performed as described in [33] and [51], respectively. ROS, phloroglucinol and aniline blue staining was quantified on micrographs using ImageJ software [76].

### JA measurements

#### Chemicals

Jasmonic acid-d_0_ and jasmonic acid-d_5_ were obtained from Santa Cruz Biotechnology, Inc. (Dallas, TX, USA). Formic acid was obtained from Merck (Darmstadt, Germany), ethyl acetate and acetonitrile (LC-MS grade) were obtained from Honeywell (Seelze, Germany). Water used for chromatographic separations was purified with an AQUA-Lab – B30 – Integrity system (AQUA-Lab, Ransbach-Baubach, Germany).

#### Sample preparation

Approximately 35 to 40 seedlings per flask were grown in liquid culture (1/2 MS, 0.3% sucrose) under continuous light and 18°C for seven days followed by treatment with mock or 600 nM isoxaben for seven hours and harvesting in liquid nitrogen. The grinded plant material (100-200 mg) was placed in a 2 mL bead beater tube (CKMix-2 mL, Bertin Technologies, Montigny-le-Bretonneux, France), filled with ceramic balls (zirconium oxide; mix beads of 1.4 mm and 2.8 mm), and an aliquot (20 *μ*L) of a solution of acetonitrile containing the internal standard (-)*trans*-jasmonic acid-d_5_ (25 *μ*g/mL), was added. After incubation for 30 min at room temperature, the tube was filled with ice-cold ethyl acetate (1 mL). After extractive grinding (3 × 20 s with 40 s breaks; 6000 rpm) using the bead beater (Precellys Homogenizer, Bertin Technologies, Montigny-le-Bretonneux, France), the supernatant was membrane filtered (0.45 *μ*m), evaporated to dryness (Christ RVC 2-25 CD *plus*, Martin Christ Gefriertrocknungsanlagen GmbH, Osterode am Harz, Germany), resolved in acetonitrile (70 *μ*L) and injected into the LC-MS/MS-system (2 *μ*L).

#### Liquid Chromatography-Triple Quadrupole Mass Spectrometry (LC-MS/MS)

Phytohormone concentrations were measured by means of UHPLC-MS/MS using a QTRAP 6500+ mass spectrometer (Sciex, Darmstadt, Germany) in the multiple reaction monitoring (MRM) mode. Positive ions were detected at an ion spray voltage at 5500 V (ESI+) and the following ion source parameters: temperature (550°C), gas 1 (55 psi), gas 2 (65 psi), curtain gas (35 psi), entrance potential (10 V) and collision activated dissociation (3 V). The temperature of the column oven was adjusted to 40°C. For plant hormone analysis, the MS/MS parameters of the compounds were tuned to achieve fragmentation of the [M+H]+ molecular ions into specific product ions: (-)trans-jasmonic acid-d_0_ 211→133 (quantifier) and 211→151 (qualifier), (-)trans-jasmonic acid-d_5_ 216→155 (quantifier) and 216→173 (qualifier). For the tuning of the mass spectrometer, solutions of the analyte and the labelled internal standard (solved in acetonitrile:water, 1:1) were introduced into the MS system by means of flow injection using a syringe pump. Separation of all samples was carried out by means of an ExionLC UHPLC (Shimadzu Europa GmbH, Duisburg, Germany), consisting of two LC pump systems (ExionLC AD), an ExionLC degasser, an ExionLC AD autosampler, an ExionLC AC column oven – 240 V and an ExionLC controller. After sample injection (2 *μ*L), chromatography was carried out on an analytical Kinetex F5 column (100 × 2.1 mm^2^, 100 Å, 1.7 *μ*m, Phenomenex, Aschaffenburg, Germany). Chromatography was performed with a flow rate of 0.4 mL/min using 0.1% formic acid in water (v/v) as solvent A and 0.1% formic acid in acetonitrile (v/v) as solvent B, and the following gradient: 0% B held for 2 min, increased in 1 min to 30% B, in 12 min to 30% B, increased in 0.5 min to 100% B, held 2 min isocratically at 100% B, decreased in 0.5 min to 0% B, held 3 min at 0% B. Data acquisition and instrumental control was performed using Analyst 1.6.3 software (Sciex, Darmstadt, Germany).

### Hypocotyl and root measurements

For measuring etiolated hypocotyl length, seedlings were grown for five days on half-strength MS agar supplemented with 0.3 % sucrose. Seedlings were photographed and hypocotyl length was measured using ImageJ. For root growth assays, seedlings were grown for seven days in long-day conditions at 21°C on half-strength MS agar supplemented with 0.3 % sucrose. Plates were inclined at 10 degrees. Root length was measures using ImageJ. For root growth recovery assays seedlings were grown on half-strength MS agar supplemented with 0.3 % sucrose. Seeds were stratified for two days followed by incubation at 21°C in long day conditions for seven days. Plates were inclined at 10 degrees. Individual seedlings were transferred to plates containing either 0.01 percent DMSO (mock) or 600 nM isoxaben for 24 hours. After treatment, seedlings were transferred onto half-strength MS agar plates supplemented with 0.3 % sucrose. The position of the root tip was marked under a dissection microscope. Root length was measured every 24 hours for up to three days. For root cell bulging assays seedlings were grown for seven days in long-day conditions at 21°C on half-strength MS agar supplemented with 0.3 % sucrose. Plates were inclined at 10 degrees. Individual seedlings were first transferred into liquid medium for two hours habituation followed by treatment with 600 nM isoxaben for up to seven hours. To take images, seedlings were stained with 4 μM FM4-64 for 10 minutes and imaged using confocal microscopy. Confocal micrographs were acquired at each time point. All hypocotyl length or root measurements were performed in double-blind fashion.

### Statistics

Statistical analysis was performed with PRISM8 software (GraphPad Software, San Diego, USA).

### Microscopy and art work

Images of seedlings exhibiting phloroglucinol staining were taken on a Leica MZ16 stereo microscope equipped with a Leica DFC320 digital camera. Images of hypocotyl and root length were taken on a Leica SAPO stereo microscope equipped with a Nikon Coolpix B500 camera. Aniline blue-stained cotyledons and root cell bulging were imaged with an Olympus FV1000 setup using an inverted IX81 stand and FluoView software (FV10-ASW version 01.04.00.09) (Olympus Europa GmbH, Hamburg, Germany) equipped with a 10x objective (NA 0.3). For assessing cell bulging a projection of a 5 μm z-stack encompassing seven individual optical sections was used. Aniline blue fluorescence was excited at 405 nm using a diode laser and detected at 425 to 525 nm. H_2_DCFDA and EGFP fluorescence excitation was done at 488 nm using a multi-line argon laser and detected at 502 to 536 nm. For the direct comparisons of fluorescence intensities, laser, pinhole and gain settings of the confocal microscope were kept identical when capturing the images from the seedlings of different treatments. Scan speeds were set at 400 Hz and line averages at between 2 and 4. Measurements on digital micrographs were done using ImageJ software [76]. Images were adjusted for color and contrast using Adobe Photoshop CC software (Adobe, San Jose, USA).

## Acknowledgments

We acknowledge Ramon Torres Ruiz and other members of the Schneitz lab for helpful discussion and suggestions. We thank Herman Höfte for *the1* alleles. We also thank Lynette Fulton, Silke Robatzek, Martin Stegmann and Sebastian Wolf for critical reading of the manuscript. We further acknowledge the support of the Center for Advanced Light Microscopy (CALM) at the TUM School of Life Sciences. This work was funded by the German Research Council (DFG) through grants SFB924 (TP B12) to CD and SFB924 (TP A2) to KS.

## AUTHOR CONTRIBUTIONS

A.C., C.D. and K.S. designed the research. A.C., X.C., J.G., B.L. and R.H. performed research. A.C., X.C., J.G., B.L., C.D. and K.S. analyzed the data. K.S. wrote the paper.

## Supporting information

**S1 Fig. *SUB* affects isoxaben-dependent induction of *CCR1* and *PDF1.2*.**

**Table S1. Primers used in this study.**

